# Real-time high-resolution microscopy reveals how single-cell lysis shapes biofilm matrix morphogenesis

**DOI:** 10.1101/2024.10.13.618105

**Authors:** Georgia R. Squyres, Dianne K. Newman

## Abstract

During development, multiscale patterning requires that cells organize their behavior in space and time. Bacteria in biofilms must similarly dynamically pattern their behavior with a simpler toolkit. Like in eukaryotes, morphogenesis of the extracellular matrix is essential for biofilm development, but how it is patterned has remained unclear. Here, we explain how the architecture of eDNA, a key matrix component, is controlled by single cell lysis events during *Pseudomonas aeruginosa* biofilm development. We extend single-cell imaging methods to capture complete biofilm development, characterizing the stages of biofilm development and visualizing eDNA matrix morphogenesis. Mapping the spatiotemporal distribution of single cell lysis events reveals that cell lysis is restricted to a specific biofilm zone. Simulations indicate that this patterning couples cell lysis to growth, more uniformly distributing eDNA throughout the biofilm. Finally, we find that patterning of cell lysis is organized by nutrient gradients that act as positioning cues.

## Introduction

Multicellularity – the assembly of individual cells into cooperative groups – confers major advantages to constituent cells^1,2^. Cells in the group can take on specialized roles, enabling them to orchestrate complex processes with greater efficiency. To achieve the benefits of multicellularity, cells within the group must pattern their behavior to perform the necessary tasks at the right place and time^3^. Organizing this patterning such that individual cells can collectively generate group-scale properties is a challenging task. Multicellular organisms have evolved elaborate cellular organizational strategies. In metazoan development, for example, a battery of signaling systems specify the activation of gene regulatory networks that determine cell fate^4^. Moreover, cells do not exist in a vacuum: multicellular assemblages are encased in extracellular materials that the cells themselves generate and whose composition and organization are patterned^5^. Yet, how the co-development of cells and their extracellular matrix environment is coordinated is only beginning to be understood.

We now know that multicellularity is not restricted to eukaryotes: most bacteria also live in multicellular collectives called biofilms^6,7^. Bacteria in biofilms must also solve the problem of organizing themselves in space and time. Indeed, studies of gene expression in biofilms have demonstrated reproducible spatiotemporal patterning^8–11^. However, the nature of the spatial and temporal patterning cues, how these cues are sensed and decoded by cells, and how this leads to community-scale function are largely unclear^12^. While important parallels exist in the cellular patterning challenges faced by bacterial biofilm and eukaryotic multicellular development, bacterial biofilm development is likely to contain distinct features. Bacterial cells are comparatively simple, and there is no overt molecular homology between bacterial and eukaryotic developmental systems. Bacteria also face unique challenges in orchestrating development: biofilms must develop across a range of environmental conditions, and bacteria must undergo development while maintaining the ability to live individually. It is therefore likely that studying bacterial biofilm organization will reveal both shared and novel mechanisms of pattern formation, making bacterial biofilms an attractive experimental system.

One of the most important processes in eukaryotic development is the construction of the extracellular matrix^13^. Similarly, the bacterial biofilm matrix is made up of polysaccharides, proteins, and extracellular DNA (eDNA), and helps define the biofilm’s physical and chemical properties and drives biofilm morphogenesis^14^. The biofilm extracellular matrix can take on complex architectures over the course of development, but how individual cells in the biofilm orchestrate matrix morphogenesis is not well understood^15^. eDNA is a major matrix component in most biofilms, and degradation of eDNA often leads to biofilm disruption^16–18^. In the pathogenic bacterium *Pseudomonas aeruginosa*, the eDNA matrix can take on a variety of architectures during biofilm development^19,20^. eDNA is released into the matrix by programmed cell death, in which a subpopulation of cells explosively lyse by expressing genes that degrade their cell wall^21^. This cell lysis must be carefully controlled to drive the morphogenesis of the eDNA matrix; however, the patterning of these lysis events during biofilm development, the underlying patterning mechanisms, and the consequences for matrix architecture are unknown. How can lysis be controlled at the single cell level to specify matrix morphogenesis at the community scale?

One of the challenges to studying the multiscale patterning of cell behavior in biofilms is that, until recently, high-resolution imaging approaches to visualize biofilm development have been limited. The study of eukaryotic development has often relied on imaging methods to visualize cell position and gene expression in the embryo, but analogous experiments in biofilms have been difficult to achieve. New methods have enabled live single-cell imaging of biofilm development, but they are generally limited in total duration and often fail in mature biofilms, a challenge for the study of complex, multi-stage processes like matrix morphogenesis^22^. In this study, we introduce a method that overcomes this technical limitation and use it to investigate the patterning of cell lysis during biofilm development, the consequences of patterned cell lysis for eDNA matrix morphogenesis, and the spatiotemporal cues used to organize this patterning. Our approaches are inspired by the study of developmental patterning in eukaryotes, and our results in turn illuminate novel features of developmental biology in bacteria. A unique feature of studying development in bacteria is that, due to their rapid growth rate, it is possible to study the full developmental life cycle from start to finish.

## Results

In designing our study, we drew inspiration from developmental biology both technically, in considering which methods were needed, and conceptually, in the experimental logic used to interrogate developmental patterning. Because cell lysis is carried out by individual cells and is expected to occur at low frequency^23,24^, we needed a method that allowed us to image biofilms with single-cell resolution. However, because cell lysis shapes the ultimate morphology of the eDNA matrix, imaging throughout the duration of biofilm development was necessary: cell lysis events that occur very early in development can impact the final architecture of the eDNA matrix several days later. Next, these images had to be analyzed quantitatively-single cell lysis events were placed onto a high-resolution map of the biofilm to determine their position, and the resulting distributions for evidence of spatiotemporal regulation were examined. After this, to characterize the impact of organized cell lysis on patterning of eDNA in the matrix, we needed to characterize the eDNA distribution over time and understand how single cell-scale lysis events could specify matrix morphogenesis at the biofilm scale. Finally, to understand how spatiotemporal regulation of cell lysis can be achieved, we needed to identify and test candidate regulatory cues. By analogy to foundational experiments in eukaryotic development, this can be done by changing the geometry of potential patterning signals and measuring for predictable changes in the localization and timing of cell lysis.

### Live, single-cell imaging of *P. aeruginosa* biofilm development

We first sought a method for live imaging of biofilm development at high spatiotemporal resolution across complete development. Previous methods of biofilm imaging that achieved multi-day duration had limited spatial and temporal resolution^25^. Recent breakthroughs in single-cell biofilm imaging achieve the necessary spatiotemporal resolution but are limited in duration^26,27^. These methods also fail in mature biofilms because fluorescent protein maturation requires oxygen, and mature biofilms contain steep oxygen gradients^28^ (Figure S1A). However, the study of matrix morphogenesis requires that we capture all stages of biofilm development across several days while maintaining single-cell resolution (Figure 1A). We began by implementing a flow cell microfluidic method to visualize biofilm growth, using a 2 cm long x 2 mm wide x 200 µm tall microfluidic growth chamber to grow flow cell biofilms of *P. aeruginosa* UCBPP-PA14 (Figure 1B). The flow cell was seeded with planktonic cells at the start of the experiment. Cells were supplied continuously with fresh media, and biofilms developed over several days. The flow cell was then mounted on a spinning disk confocal for live, 3D time-lapse imaging. To overcome the challenges of fluorescent protein-based approaches, we combined this with a “fluorescence exclusion” approach by adding 20 µM fluorescein to the flow media^29^. Fluorescein is not cell permeable, meaning that cells appear dark against a fluorescent background. These images can be inverted to look like standard images of internal cellular fluorescence and provide single-cell resolution (Figure 1B, S1B). Fluorescein is a practical choice for these experiments because it is inexpensive and non-toxic^30^, and we confirmed that the presence of 20 µm fluorescein did not affect *P. aeruginosa* growth (Figure S1C). Most importantly, because the fluorescein is replenished continuously, there is no detectable photobleaching (Figure S1A). This allowed us to extend single-cell biofilm imaging methods to capture all stages of biofilm development over a week or more. These long duration experiments are essential for the study of biofilm development: they allow single cell behavior in early development to be connected to their ultimate developmental outcomes.

**Figure 1:**
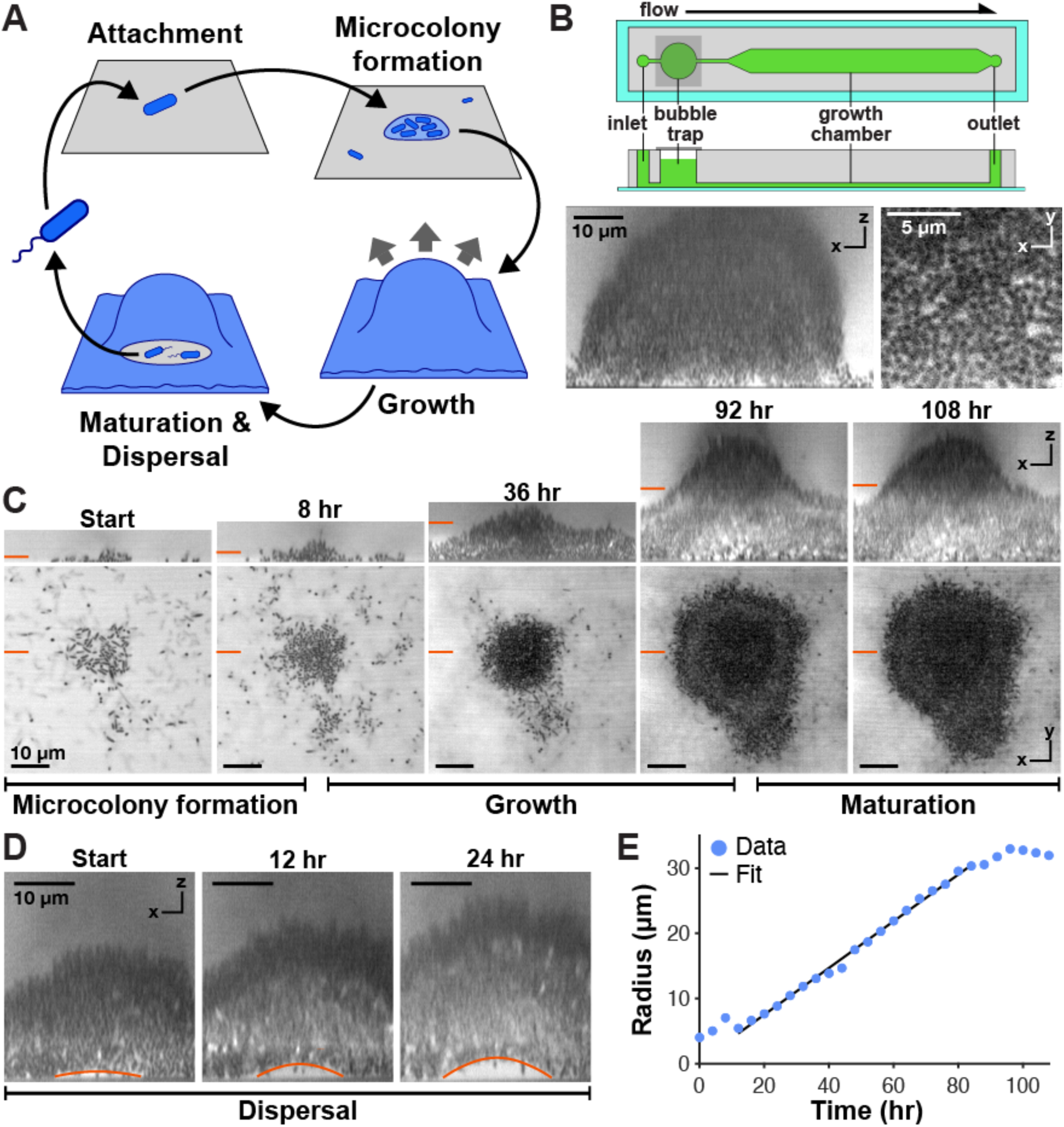
Live, single-cell imaging of biofilm development in *P. aeruginosa*. **(A)** The standard model of *P. aeruginosa* biofilm development^55^. **(B)** Biofilm imaging at single-cell resolution using fluorescence exclusion microscopy, in which fluorescein is added to the medium and used as a negative stain. Top: Schematic of the flow cell microfluidic device used in this study. Bottom: Confocal XZ (left) and XY slices of single cells (right) in a *P. aeruginosa* biofilm visualized by fluorescence exclusion imaging. **(C)** Live, multi-day imaging of all stages of biofilm development. XZ (top) and XY (bottom) sections from 4D confocal imaging data. Orange markers indicate the position of the orthogonal section. See also Video S1. **(D)** Hollow core formation during seeding dispersal is caused by biofilm deformation: the biofilm core lifts off the surface. Confocal XZ section, orange line indicates the bottom surface of the biofilm that separates from the surface and moves upward. See also Video S2. **(E)** Biofilm growth over time. Biofilm radius increases linearly during the growth phase (black line: linear fit).

With this method, we imaged *P. aeruginosa* flow cell biofilm development. We captured the key stages of biofilm development: cell attachment and microcolony formation, growth, maturation, and dispersal (Figures 1C and 1D, Video S1). Initially, cells were present at low density, moving along the surface by twitching motility (Figure S2A). Over time, microcolonies emerged; this process typically took 3 days, with some variation between replicates. These colonies then grew over the following days into large, hemispherical biofilms, like the biofilm architecture that is often seen in microfluidic flow cells^24^ (Figure 1C). During this time, cells in the biofilm core become tightly compacted, vertically aligned, and largely immobile; only cells at the biofilm periphery exhibit growth and motility (Figure S2D, Video S1). The biofilm grows with linearly increasing radius, consistent with cell growth that is localized to the outer surface, as in flow cell biofilms of *V. cholera*^27^ (Figure 1E). After a growth period, biofilm growth halts, associated with the transition into the maturation phase (Figures 1E and 1F). In a subpopulation of biofilms, maturation is followed by the appearance of open cavities in the biofilm core containing motile cells, which is when the dispersal process commences that ends the *P. aeruginosa* biofilm life cycle^31^ (Figures 1F, S2D). Unexpectedly, we observed that this cavity forms by the biofilm detaching from the surface and lifting upward, in contrast to previous models that suggested that the hollow core forms by cell death or by departure of motile cells^32,33^; this type of mechanical event is very difficult to detect without continuous live imaging (Figure 1D, Video S2). In summary, our system allows us to capture major features of *P. aeruginosa* biofilm development, while increased spatial and temporal resolution provides novel insight.

### eDNA matrix architecture during biofilm development

Next, we sought to visualize eDNA matrix morphogenesis throughout development. We added a cell-impermeant eDNA stain, either DiTO-1 or propidium iodide (PI) to the medium at low concentrations; all experiments were repeated with both stains to verify that the results did not depend on the specific stain used. This allowed us to monitor the structure of the eDNA matrix in biofilms over time (Figures 2A and 2B). During the initial attachment phase, cell lysis events lead to a dense coating of eDNA on the chamber surface (Figure S2B). This eDNA has been shown to promote adhesion of *P. aeruginosa* biofilms to the surface during the early stages of biofilm development^16^. As microcolonies form, they are dense with eDNA, consistent with previous studies^23^ (Figure 2A). At this stage, cells in microcolonies are individually enmeshed in eDNA (Figure S2C). Next, during growth, punctate foci of eDNA appear throughout the biofilm (Figure 2A, Video S3). At this stage, cells in the core of the biofilm are essentially immobile, and similarly, these eDNA foci do not substantially move or spread from their initial location once they appear. The dense eDNA layer deposited during the initial attachment and microcolony stages remains present at the base of the biofilm. Some lysis continues in this region, particularly during the maturation phase, consistent with previous reports of cell lysis in the biofilm core as a precursor to biofilm dispersal^33,34^. Finally, at the end of biofilm growth, this dense basal layer of eDNA moves upward to create the hollow cores associated with dispersal (Figure 2B, Video S2). In summary, different eDNA matrix morphologies characterize different stages of biofilm development. Once deposited, the eDNA is largely stationary, with only minimal movements until the dispersal stage and no overt loss of eDNA in the matrix over time. This means that the morphogenesis of the eDNA matrix is specified entirely by the location of cell lysis events.

**Figure 2:**
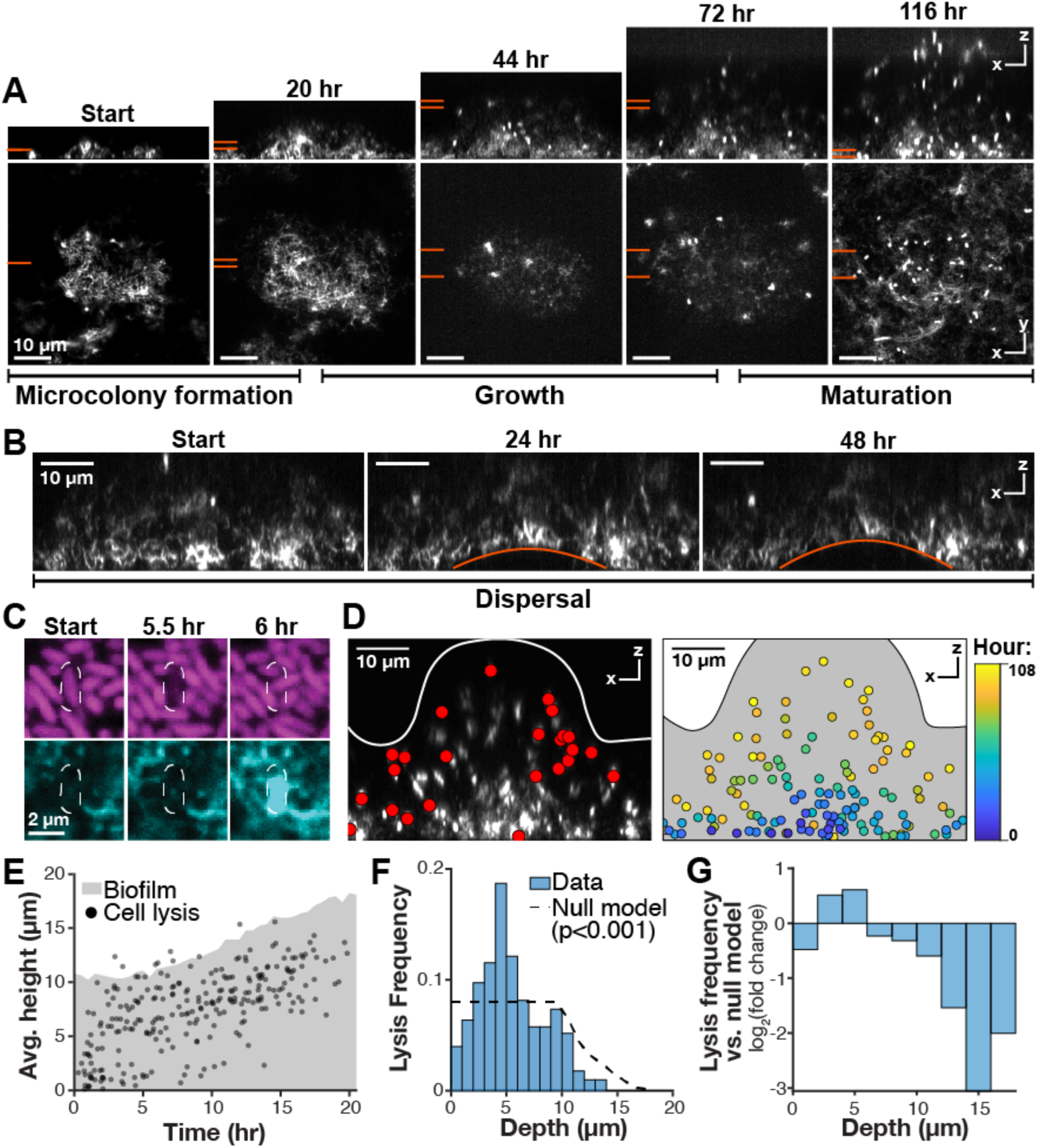
Cell lysis is patterned during *P. aeruginosa* biofilm growth. **(A)** Morphogenesis of the eDNA matrix during biofilm development. During the microcolony phase, eDNA forms a dense mesh. During growth, individual foci of eDNA appear in the growing region. During maturation, lysis begins to occur in the biofilm core region, which previous research has suggested facilitates dispersal^33^. XZ and XY maximum intensity projected views from 4D confocal imaging data, eDNA stained with DiTO-1. Orange bars: region used for maximum intensity projected region in the orthogonal view. **(B)** The eDNA matrix detaches from the surface and moves upward during biofilm dispersal. The hollow biofilm core is visible at the bottom of the image. Confocal XZ section, eDNA stained with DiTO-1. Orange line indicates the bottom surface of the biofilm that separates from the surface and moves upward. See also Video S2. **(C)** Microscopy method for detecting single cell lysis events. Cell lysis causes the disappearance of the cell accompanied by the appearance of a labeled focus of eDNA. Confocal biofilm section, magenta: mApple cytoplasmic fluorescence, cyan: DiTO-1. **(D)** Localization of single cell lysis events during biofilm growth. Left: Recent cell lysis events appear in a specific biofilm region. Confocal XZ slice of eDNA in a representative biofilm labeled with PI. Overlaid red dots indicate cells that lysed during the previous 8 hours, white line indicates biofilm surface after 8 hours. Right: Location of all cell lysis events during the growth phase, color-coded by time. Lysis events were mapped by quantitative image analysis. Gray region indicates the final shape of the biofilm. See also Video S3. **(E)** Cell lysis occurs in a specific region that follows the growing edge of the biofilm. Representative time course of cell lysis during biofilm development, from 6 biofilms across 3 biological replicates. Gray region: average biofilm height, and black points: cell lysis events. **(F)** Cell lysis is spatiotemporally regulated in growing biofilms. Blue bars: integrated distribution of cell lysis events as a function of depth, *i.e.* distance from the lysed cell to the closest point in the biofilm surface as the biofilm grows. Black dashed line: null model in which cell lysis is not depth-dependent, obtained by simulation. These distributions are significantly different (p<0.001) by Mann-Whitney U test. **(G)** Comparison of lysis frequency to null model. Lysis occurs more often than expected in a region approximately 5 µm below the biofilm surface, and less often elsewhere. Log2(fold change) of lysis depth distribution versus simulated null model.

### Patterned cell lysis specifies biofilm matrix morphogenesis

To understand how single cell lysis events lead to eDNA matrix morphogenesis, we next analyzed our time lapse microscopy data to identify single cell lysis events and map their location in space and time (Figures 2C-2E). Because of the use of cell-impermeant DNA stains, cell lysis is accompanied by the appearance of a fluorescently labeled focus of eDNA^23^ (Figure 2C). We tracked the appearance of these foci to map cell lysis. Cell lysis occurs throughout development, but as expected, lysis is restricted to a very small subpopulation: approximately 1 in 10,000 cells lyses during biofilm growth. Nearest-neighbor analysis indicates that locally, the distribution of cell lysis events is stochastic (Figure S3A). At larger scales, however, cell lysis is spatially and temporally patterned during the biofilm growth phase. Specifically, we observed that cells predominantly lyse in a layer at intermediate depth within the biofilm (neither at the biofilm core nor the extreme periphery) that moves with the biofilm as it grows. Analysis confirmed that cell lysis predominantly occurs in a region centered 5 µm below the biofilm surface (Figures 2F and 2G). This patterning was not due to limitations in eDNA dye labeling, and did not depend on the specific eDNA dye used (Figures S1D-E and S3B). We used simulation to construct a null model: the predicted cell lysis distribution in the absence of spatiotemporal regulation (see “Simulation and Modeling” in Methods for details). This cell lysis distribution is significantly different from the unregulated null model (Figure 2F and 2G), indicating that cell lysis during *P. aeruginosa* biofilm growth is spatiotemporally patterned.

Single cell lysis events occur quickly in individual cells, but morphogenesis of the eDNA matrix occurs over several days and across the whole biofilm. How does cell lysis patterning shape the biofilm matrix at a larger scale? To understand this relationship, we constructed two models, described in “Simulation and Modeling” in Methods (Figure 3A). In one, cell lysis is uniformly distributed throughout the biofilm, such that every cell has an equal probability of lysing. In the other, the distribution of cell lysis is patterned to match the one we observed experimentally, occurring at 5 µm below the biofilm surface. For both models, the growth dynamics were matched to those we observe experimentally. Comparing these models yielded specific predictions that we compared to our experimental results. First, we examined the consequences of patterned lysis on the amount of eDNA released into the matrix during biofilm development. In the patterned model, we found that cell lysis is matched to biofilm growth, meaning that the total amount of eDNA contributed to the matrix scales with the total biofilm volume (Figure 3B). Our experimental data reflects this coupling (Figure 3B). This does not happen in the uniform model; instead, the amount of eDNA in the biofilm increases faster than the biofilm grows (Figure 3B). Second, we investigated what impact regulated cell lysis has on the final structure of the eDNA matrix. Uniquely in the case where cell lysis is patterned, the final eDNA matrix distribution is constant except for the first few microns of the biofilm that are depleted for eDNA (Figures 3C and 3D). In the uniform model, the distribution of eDNA is highest at the biofilm core and decreases towards the periphery. We measured the final distribution of eDNA in our biofilms and confirmed that it matched the predicted distribution from our simulation (Figures 3C and 3D). Together, these results indicate that spatiotemporal regulation of cell lysis leads to coupling between cell lysis and biofilm growth, resulting in a uniform eDNA matrix architecture at the biofilm scale.

**Figure 3:**
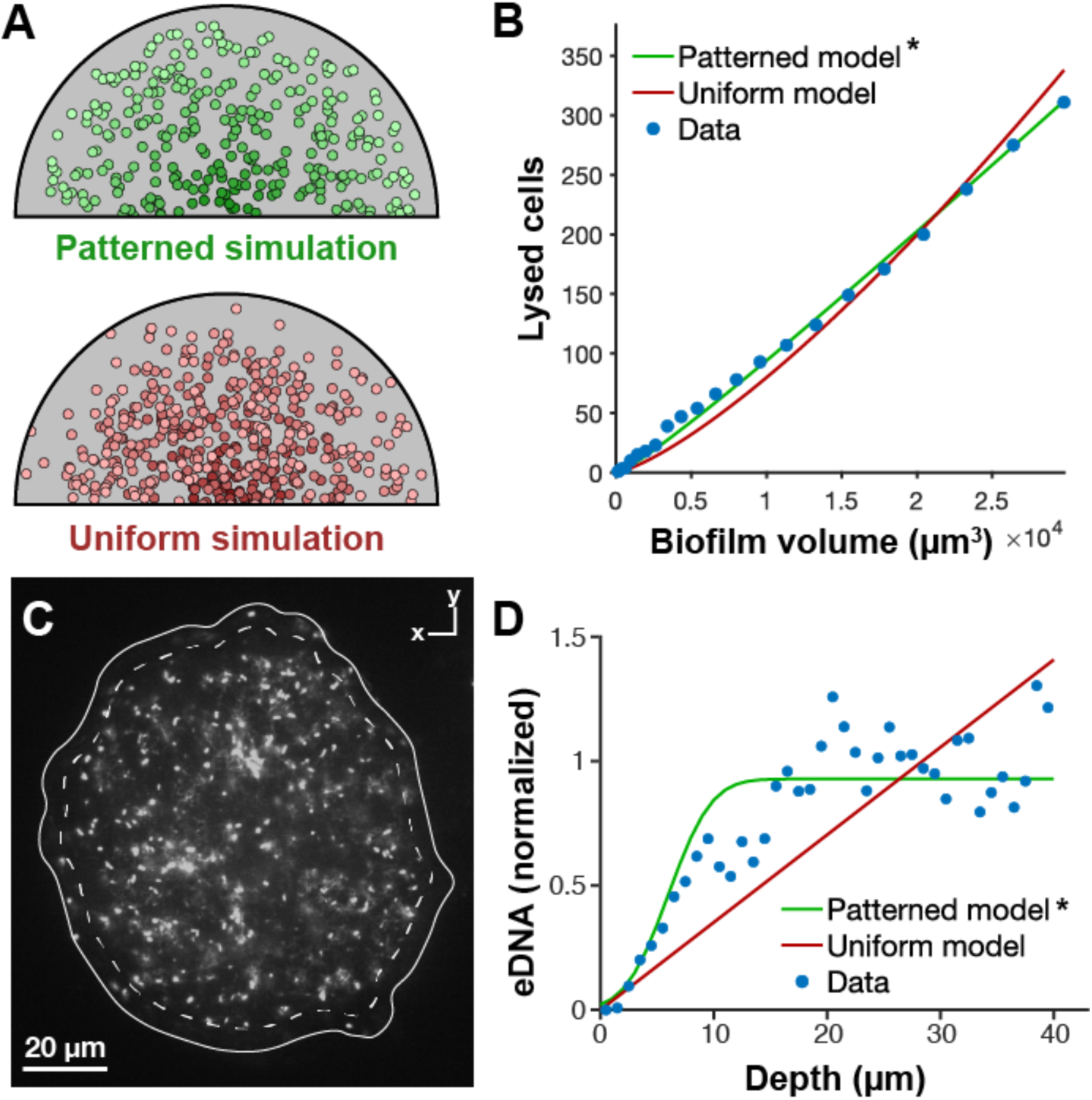
Patterned cell lysis specifies the amount and distribution of eDNA in the biofilm matrix. **(A)** Simulations of different modes of cell lysis regulation integrating cell lysis events over time. In the patterned simulation (top), the cell lysis distribution is matched to what we observe over biofilm development, with a peak 5 µm below the biofilm surface. In the uniform simulation (bottom), cells lyse with uniform probability everywhere. See “Simulations and modeling” in Methods. Visualization of simulation results: gray hemisphere indicates the final biofilm contour, and points indicate lysis events; points are colored dark to light in time. **(B)** Patterned lysis leads to coupling between biofilm growth and cell lysis. In the patterned simulation (green) the total number of lysed cells is directly proportional to the total biofilm volume; in the uniform model (red) this is not the case. Lysis data (blue points) reflects this proportionality. *Best fit, meaning the model that maximized R^2^ after fitting, see Methods for details. **(C)** The biofilm eDNA distribution: eDNA is depleted from the first few microns of depth but is otherwise largely uniform. Representative confocal section from 3 biological replicates, eDNA stained with PI, white line indicates biofilm surface, dashed line indicates 5 µm depth. a**(D)** Patterned cell lysis specifies the eDNA matrix distribution. In the patterned simulation, eDNA is spatially uniform throughout the biofilm except at the periphery, and eDNA distributions measured by imaging show a similar distribution. In the uniform simulation the eDNA concentration instead scales with depth. *Best fit.

### Exogenous nutrient gradients pattern cell lysis

Finally, we considered what cues may lead to patterned cell lysis: why do cells die in a band 5 µm beneath the biofilm surface? Candidate cues for this patterning must satisfy several criteria. To encode position, the cue must vary with biofilm depth, and its depth profile must move with the growing biofilm surface. It must vary on a 5 µm length scale, and cells must be able to sense it and respond accordingly. We hypothesized that gradients of exogenous nutrients (e.g. carbon and oxygen) might serve as these cues. Biofilms quickly consume nutrients from their environment, leading to the formation of steep gradients where the concentrations of nutrients decrease with increasing depth into the biofilm^35^. As a result, biofilms exhibit complex, reproducible patterning of metabolism and other processes that depend on local nutrient concentration such as sporulation and dormancy ^36–38^. We therefore wondered whether in this case a non-metabolic process might take advantage of these gradients to establish depth-based patterning.

In *P. aeruginosa*, nutrient depletion alone is insufficient to drive cell lysis: *P. aeruginosa* cells do not lyse even after several days of severe energy limitation, and explosive cell lysis requires active expression of lysis-causing genes^21^. However, activation of the cell lysis pathway in *P. aeruginosa* can depend on the environment, and conditions where oxygen is present but cells are limited for carbon are known to be particularly toxic, and we verified in stationary phase liquid culture experiments that more eDNA is released in this condition^39^ (Figure 4A). Because the carbon concentration in these experiments is low (the medium contains 300 µM glucose; *P. aeruginosa* flow cell biofilms are commonly grown under low carbon concentrations^20^), we reasoned that cell lysis might be occurring in a biofilm region where carbon is limited but oxygen is present. In this model, cells do not die at the biofilm periphery because sufficient carbon is present, and do not die at the core because oxygen has been depleted (Figure 4B).

**Figure 4:**
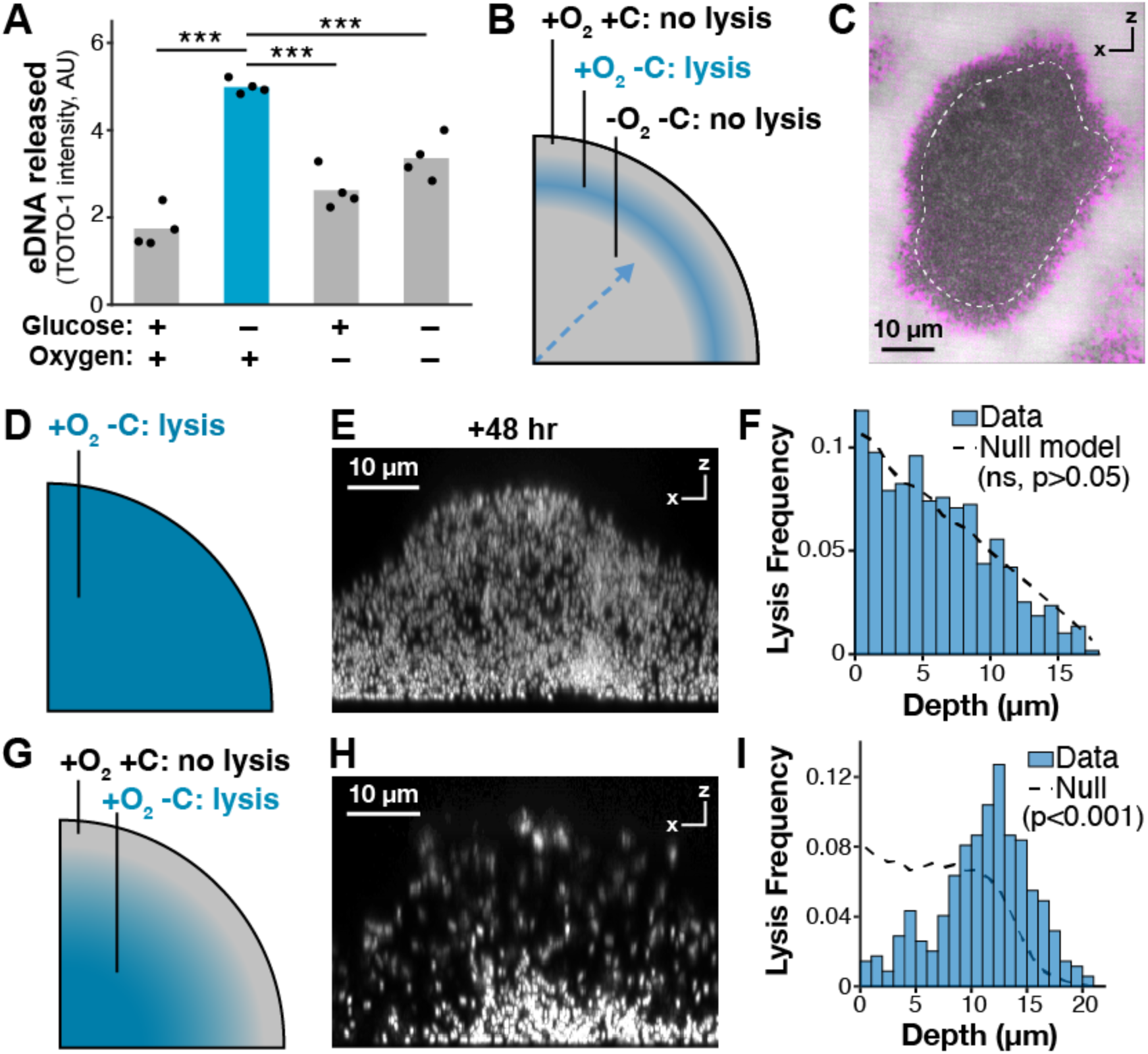
Cell lysis in biofilms is patterned by dynamic nutrient gradients. **(A)** Cell lysis occurs preferentially in low-carbon, high-oxygen conditions. Stationary phase cultures of *P. aeruginosa* were incubated in the indicated conditions for 48 hours, then stained with TOTO-1 to quantify eDNA release. Points: individual replicates, bars: mean of replicates. Blue bar indicates predicted lysis condition. ***p<0.001 by T-test. **(B)** Model for cell lysis regulation in biofilms. Biofilms contain steep nutrient gradients, meaning that nutrient conditions can vary sharply with depth, and move in time as space as the biofilm develops (dashed blue line); cell lysis occurs in a region of the biofilm containing low oxygen and high carbon. **(C)** *P. aeruginosa* biofilms contain steep nutrient gradients. Confocal cross-section of a *P. aeruginosa* biofilm; regions labeled magenta are areas of the biofilm that exhibited cell growth and motility during the previous 4 hours. Cell lysis occurs preferentially beneath this growing region (white dashed lines indicates contour 5 µm below the biofilm surface). **(D-F)** Cell lysis patterning is lost in the absence of dynamic nutrient gradients. D: After growing biofilms to the middle of their growth phase, carbon was removed from the flow medium, leading to high oxygen and low carbon throughout E: After 48 hours of no-carbon media, cell lysis is visible throughout the biofilm. Confocal XZ section, lysis visualized with PI. See also Video S4. F: Cell lysis frequency is no longer spatiotemporally patterned in the absence of dynamic nutrient gradients. Lysis distribution and null model determined as in Figure 2F; these distributions are not significantly different (p>0.05) by Mann-Whitney U test. **(G-I)** Increasing oxygen drives increased lysis in the biofilm core. G: In a more oxygen-replete region of the flow cell, oxygen concentration throughout the biofilm is increased. H: eDNA distribution in an oxic biofilm, indicating increased lysis near the biofilm core. Confocal XZ section, lysis visualized with PI. See also Video S5. I: Cell lysis distribution: cell lysis is enhanced in the biofilm core when oxygen is present.

We evaluated this hypothesis in several ways. First, we compared the localization of cell lysis in the biofilm to the location of cell growth. If nutrient gradients are indeed formed in these biofilms, we expected to find that cell growth is restricted to the outer layers of the biofilm where nutrient levels are highest, as has been reported for other flow cell biofilms^27^. Because we can visualize individual cells throughout biofilm development, we can compare their positions over time to determine in which regions of the biofilm cells are rearranging by growth. By analyzing single cell biofilm time lapses, we found that during the biofilm growth phase, only cells at the outermost periphery of the biofilm are actively growing. Cell lysis is maximized at a depth of 5 µm, where cell growth no longer occurs (Figure 4C).

Next, we conducted two experiments to alter these gradients and determine the effects on cell lysis patterning (Figures 4D-4I). In each case, we grew biofilms until they were in the middle of the growth phase, and then changed the conditions to alter the shape of the nutrient gradient and observe the effects on cell lysis patterning. In the first experiment, we removed the carbon source from the medium. This leads to the biofilm becoming carbon limited throughout. We additionally expected that the oxygen gradient would collapse as cellular respiration rates decreased, leading to high oxygen everywhere. Thus, after carbon source removal we expected that the biofilm would be globally low in carbon and high in oxygen, promoting cell lysis (Figure 4D). Indeed, we found that after removing carbon from the medium, cells lysed throughout the biofilm with no spatial patterning (Figures 4E and 4F, Video S4). This effect was biofilm-specific: removal of carbon from the medium did not increase cell lysis at the single cell stage prior to biofilm formation (Figure S3D).

In the second experiment, we imaged biofilms growing near the edges of the flow chamber. Our flow cells are made of polydimethylsiloxane (PDMS), which is a gas-permeable resin. The oxygen concentration is therefore highest at the chamber edges, where oxygen can enter. We verified this by confirming that fluorescent protein maturation depended on distance from the chamber edge, indicating that more oxygen is available in this region (Figure S3C). In this case, we expected to see increased lysis at greater biofilm depth, since the deeper biofilm regions now were expected to contain oxygen but still be depleted for carbon. Indeed, we observed that cells began to lyse in the biofilm core when biofilms were more oxygenated (Figure 4G, Video S5). Together, these experiments indicate that gradients of nutrients and oxygen pattern cell lysis in *P. aeruginosa* biofilms.

## Discussion

How multiscale coordination between individual cells and larger matrix morphogenesis is achieved is an open question in many developmental systems^40^. Here, we investigated how the eDNA matrix component of bacterial biofilms is shaped by patterned lysis at the single-cell level. By characterizing the progression of eDNA matrix architecture and the tandem distribution of single cell lysis events under different experimental conditions, we discovered that eDNA deposition is matched with biofilm growth in a specific manner to yield a uniform matrix architecture. We found that biofilm cells exploit the information encoded in nutrient gradients to determine their position in the biofilm and regulate their behavior accordingly. Together, these results demonstrate that bacteria in biofilms can use self-produced nutrient gradients to pattern their behavior in space and time, leading to organized behavior at the community scale.

In eukaryotes, programmed cell death is carefully regulated to shape tissue morphogenesis throughout development^41^. Here, we have found that patterned lysis during biofilm development shape the morphogenesis of the eDNA matrix, resulting in a largely spatially uniform distribution of matrix eDNA. What impact might this matrix architecture have on biofilm fitness? eDNA has several functions in the biofilm matrix that could benefit from a uniform spatial distribution. eDNA largely specifies the mechanical properties of *P. aeruginosa* biofilms, and variation in matrix concentration across the biofilm will alter these properties^42,43^. eDNA in biofilms also serves to bind and retain small molecules involved in signaling and metabolism, and organized cell lysis may ensure that adequate eDNA is present throughout the biofilm for this purpose^44^. Organized cell lysis during growth also leads to a very low concentration of eDNA at the biofilm periphery, perhaps ensuring that bacteria in this region can freely undergo growth and motility. Matching cell lysis to growth can ensure that new eDNA is deposited in the right amount where and when it is needed, while preventing the unnecessary fitness cost of excess cell lysis in the biofilm core.

We have found that cell lysis is spatially organized in depth by coupling to specific nutrient conditions. How might this coupling be accomplished: what drives lysis specifically in high-oxygen, low-carbon conditions? We predict that nutrient information is integrated into existing pathways for *P. aeruginosa* programmed cell death. eDNA release in *P. aeruginosa* can be caused by the self-produced secondary metabolite pyocyanin, which is abundant in the oxic region of biofilms and whose autotoxicity is maximized in high-oxygen, low-carbon conditions^39,45^. Because the effect of these molecules on the cell depends on the local nutrient context, biofilm bacteria can use them to decode their position in the nutrient gradient. Additionally, pyocyanin production is quorum regulated, and quorum sensing has been implicated in regulation of *P. aeruginosa* programmed cell death^46,47^. Quorum sensing may therefore restrict the activation of this pathway to the community context where programmed cell lysis can confer a group benefit.

Here we consider a specific example of biofilm patterning in *P. aeruginosa*, but patterning in biofilms occurs broadly across processes and microorganisms. Nutrient gradients are formed ubiquitously in biofilms due to the high cell density associated with the biofilm lifestyle and the fact that consumption of substrates typically outpaces their diffusion^35^. A role for nutrient gradients as biofilm patterning cues has been considered for some time in the field^48^, and it has been suggested that these gradients may serve as morphogens to pattern biofilm development^36,49^. How morphogen distribution is controlled to pattern development is an active area of research in eukaryotic developmental biology^50^, but this research has largely focused on endogenous factors (*i.e.* proteins or small molecules that are generated by the cells). In this case, exogenous nutrient gradients are serving a morphogen-like function, in which cells determine their position in the biofilm based on their position in the nutrient gradient that is shaped by local nutrient consumption. Here, the source of the morphogen is at the growing surface of the biofilm, which allows cells to pattern their behavior based on their position relative to the leading edge, and the outcome is controlled morphogenesis of the biofilm matrix. The use of a morphogen to mark the leading edge of a growing tissue to coordinate growth with morphogenesis has parallels in other developmental systems. For example, in plant root development, a high concentration of auxin is maintained at the growing root tip, and the resulting gradient patterns the plant root as it grows^51^. In both cases, the result is continuous development of a uniform structure during growth.

What are the implications of using nutrients as morphogens, rather than the signaling molecules often used in eukaryotes? Nutrients are intriguing candidates for patterning cues because they are integrate information about the local environment, including nutrient availability and the presence of other organisms. Using nutrients as signals may allow biofilms to adapt their developmental trajectories to their environment. Because biofilms must develop in a variety of conditions, and because cells must ultimately leave the biofilm at the end of their life cycle, this nutrient information may be crucial for biofilm decision-making. Indeed, nutrient conditions shape biofilm morphogenesis in *P. aeruginosa*, and nutrients shifts can trigger biofilm dispersal over rapid timescales^52–54^. Bacteria may additionally integrate nutrient information with other environmental cues – for example, high cell density that is characteristic of biofilms – to ensure that target pathways are activated in the appropriate context. More generally, the use of nutrient gradients in biofilm patterning illustrates how bacteria may adapt the patterning rules used in eukaryotes to meet the specific challenges of biofilm development for organisms that are capable of both a solitary and multicellular lifestyle. The methods presented here allow us to visualize patterning at the cellular scale and link it to function at the biofilm scale, enabling continuing investigation of the principles that organize microbial development.

## Acknowledgements

We thank Alice Cont, Alexander Persat, and Reinaldo Alcalde for microfluidic fabrication advice, and all members of the Newman lab for helpful discussions. We gratefully acknowledge the critical support and infrastructure provided for this work by the Kavli Nanoscience Institute at Caltech, where microfluidic fabrication was conducted. This work was supported in part by the Resnick Sustainability Institute: imaging was primarily performed at the Resnick Ecology and Biosphere Engineering Facility at Caltech. Supplemental imaging was conducted at the Caltech Biological Imaging Facility. GRS is a National Mah Jongg League Fellow of the Damon Runyon Cancer Research Foundation (DRG 2439-21). This research was supported by NIH grant (2R01AI127850-06A1) to DKN.

## Author contributions

Conceptualization and Methodology: GRS and DKN; Investigation, Validation, Formal Analysis, Visualization and Software: GRS; Writing – Original Draft, Review & Editing: GRS and DKN; Supervision and Funding Acquisition: DKN.

## Methods

### Strains and strain construction

This study was conducted in *P. aeruginosa* UCBPP-PA14. We used two strains: wild type (WT), and a strain (DKN3010) constitutively expressing mApple under the control of the P_A1/04/03_ constitutive promoter^58^. To construct DKN3010, we first made a plasmid (DKN3008) for homologous recombination into the attB site, a neutral locus commonly used in *P. aeruginosa* cloning^59^. We used the plasmid pMQ30 for markerless allelic exchange^56^. We PCR-amplified two regions immediately upstream and downstream of the attB site using purified WT *P. aeruginosa* genomic DNA as template (see Table 1 for primers). We, linearized by SmaI digestion, and added the attB homology regions by Gibson assembly. *E. coli* DH10β was used as a host strain for cloning. We added a SmaI cut site in between the two attB homology regions for future cloning. Next, we inserted PA1/04/03-mApple into this plasmid to create DKN3009. We ordered a codon-optimized mApple gene fragment (Twist) and used plasmid DKN238 as the source for PA1/04/03. DKN3008 was linearized by SmaI digestion and fragments were combined by Gibson assembly. To transform the construct into *P. aeruginosa*, DNK3009 was transformed into the *E. coli* donor strain S17-1λpir, and the plasmid was moved into WT *P. aeruginosa* by biparental conjugation^60^. Following counterselection, we were left with *P. aeruginosa* strain DKN3010, which contains a markerless PA1/04/03-mApple chromosomally integrated into the attB site. Strains were verified by diagnostic PCR and sequencing.

**Table 1:**
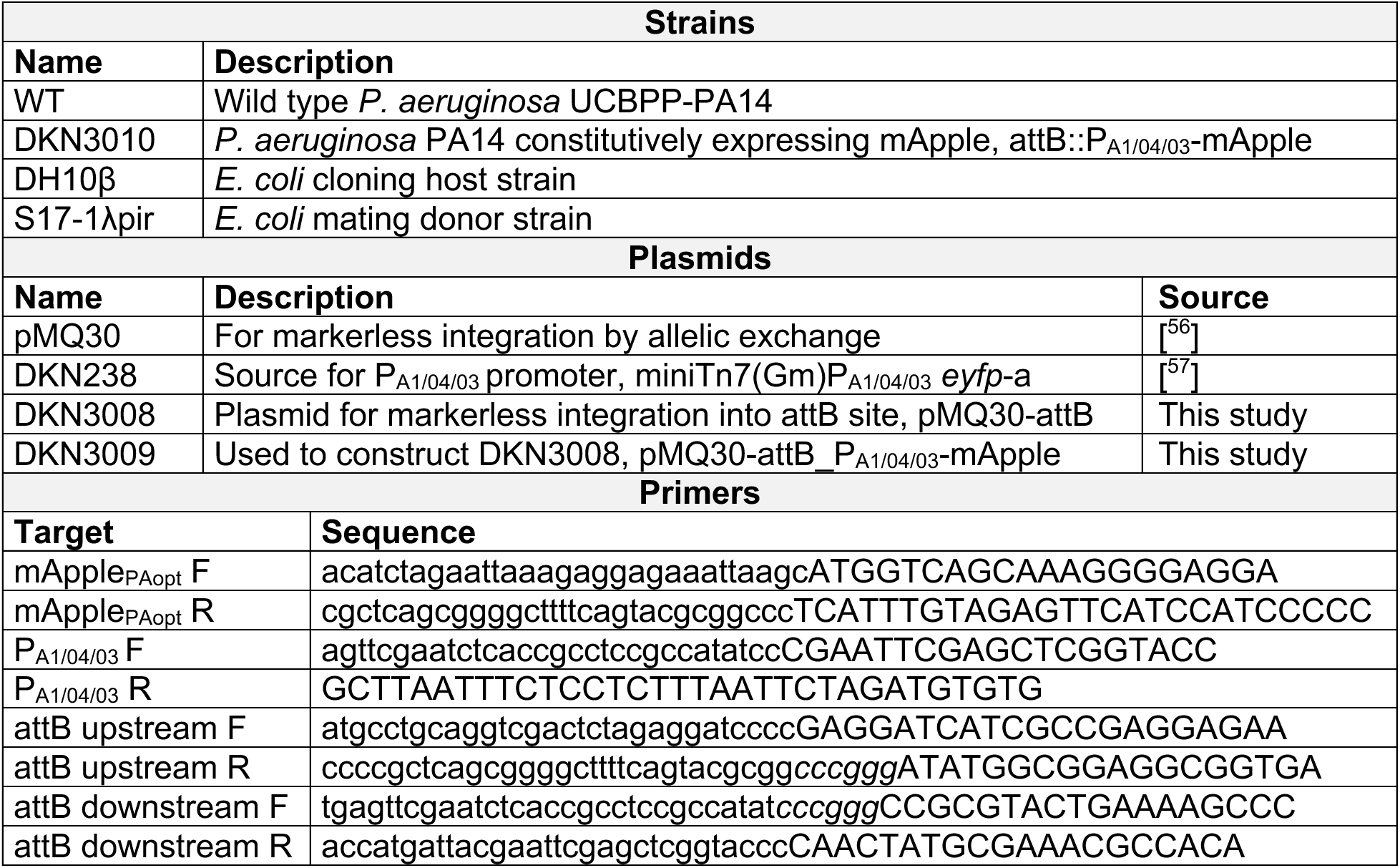
Strains, primers, and plasmids used in this study. In primer sequences, capital letters are the annealing region, lowercase letters are homologous regions for Gibson cloning, and italics indicate restriction sites.

### Culture conditions and media

Prior to each experiment, strains were streaked from cryopreserved glycerol stocks onto LB agar plates and grown overnight at 37°C. To grow starter cultures, single colonies were inoculated into 3 mL Jensen’s medium^20^ containing 70 mM glucose, 65.6 mM NaCl, 14.4 mM potassium phosphate (7.72 mM K_2_HPO_4_ + 6.67 mM KH_2_PO_4_), 95 mM sodium glutamate, 24 mM valine, 8 mM phenylalanine, 1.33 mM MgSO_4_, 0.14 mM CaCl_2_, and an EDTA-chelated trace metals mix consisting of 10 µM FeCl_3_^+^, 120 nM MnCl_2_, 20 nM CuSO_4_, 50 nM CoCl_2_, 100 nM Na_2_MoO_4_, 10 nM Na_2_SeO_3_, and 80 nM ZnSO_4_ with 100 µM Na_2_EDTA. Cultures were grown with shaking overnight at 37°C to stationary phase. For flow cell experiments, modified Jensen’s minimal medium (MJMM)^20^, a glucose minimal medium, was used. MJMM has the same composition as Jensen’s medium except that the glucose concentration was reduced to 300 µM, the amino acids were removed, and 15.1 mM (NH_4_)_2_SO_4_ was added. For carbon removal experiments, MJMM medium was used with no glucose added.

### Growth curves

For growth curves, strains were grown overnight in Jensen’s medium as described above. In the morning, they were diluted to an approximate starting OD_600_ of 0.05 into Jensen’s medium with stains added in a 96-well plate. Stains were added in the same combinations used for the biofilm imaging experiments and at the same concentrations (fluorescein: 20 µM, DiTO-1: 50 nM, PI: 500 nM). The plate was loaded into a plate reader and shaken at 37°C for 20 hours, with OD_600_ measurements taken every 10 minutes. The experiment was performed in biological quadruplicate. To compute maximal growth rates, the early portion of the blank-subtracted curve was fitted to a Gompertz function using custom MATLAB code.

### eDNA measurements in liquid culture

WT cells were grown overnight to stationary phase in Jensen’s medium in quadruplicate. In the morning, each culture was divided into four batches, with one batch assigned to each growth condition. The carbon-containing condition used MJMM medium containing 70 mM glucose, and the non-carbon condition used MJMM medium with no glucose added. For anoxic conditions, cells were moved into a vinyl anaerobic chamber (Coy), and the medium used was pre-incubated in the anaerobic chamber for several days to remove oxygen. Cells were washed once and resuspended in the appropriate medium but were not diluted: cell density remained high. Cultures were incubated for two days with shaking at 37°C. Afterward, 1 µM DiTO-1 was added to each well to stain eDNA, and plates were incubated at 4°C during staining. DiTO-1 fluorescence was measured using a fluorescence plate reader (Tecan).

### Microfluidic design

Microfluidic device design was based on a template kindly shared with us by Dr. Alexander Persat^61^. The device was made of polydimethylsiloxane (PDMS) (Sylgard 184), with the bottom made of 24x50 mm no. 1.5 optical cover glass. The cell growth chamber was 2 cm long x 2 mm wide x 200 µm tall. The growth chamber was fed by a 7 mm long x 300 µm wide x 200 µm tall inlet channel which contains a bubble trap. Tubing was connected at the start of the inlet channel and the end of the growth chamber. Device schematics are shown in Figure 1B.

### Microfluidic flow cell fabrication

Microfluidic flow cells were fabricated by conventional soft lithography. A mylar photomask of the flow cell design was ordered from FineLine Imaging, containing 12 devices arranged to fit on a 4” silicon wafer. All fabrication steps were performed in the Kavli Nanoscience Institute clean room at the California Institute of Technology. A 4” silicon wafer was cleaned, spin coated with SU-8 TK photoresist to a thickness of 200 µm following the manufacturer’s instructions, then baked on a hot plate, first for 10 minutes at 65°C followed by 80 minutes at 95°C. The photomask was applied, and the photoresist was crosslinked by flood exposure to 365 nm UV light at 15 mW/cm^2^ for 30 seconds. The wafer was baked again at 65°C for 5 minutes and 95°C for 25 minutes. Uncrosslinked photoresist was removed by 20 minutes of immersion in SU-8 developer. The wafer was rinsed and then baked at 200°C for 15 minutes. This created the final negative mold. Final height of the features was verified to be approximately 200 µm by light microscopy.

To create individual flow cells, the mold was placed into a Petri dish lined with aluminum foil folded up around the sides of the wafer to create a well into which PDMS could be poured. PDMS components were mixed in a 10:1 weight ratio and degassed in a benchtop vacuum chamber. 25 g of PDMS was poured into the mold; this was degassed again to remove bubbles introduced during pouring and then cured for 20 hours on a leveled surface in a 55°C oven. In the clean room, the cured PDMS was peeled away from the mold. Individual flow cells were cut apart and excess PDMS was cut away using a razor blade. Using a 1 mm biopsy punch, holes were punched at the start of the inlet channel and the end of the growth chamber for tubing connections; a 3 mm punch was used to punch a bubble trap hole in the middle of the inlet channel. The PDMS flow cells and 24x50 mm coverslips were rinsed with isopropyl alcohol and dried with an air gun. Immediately before use, the PDMS and glass were treated with air plasma for 1 minute in a plasma cleaner (Harrick) set to high RF power. The treated pieces were immediately pressed together to bond them, forming the complete flow cell.

### Flow cell biofilms

Cells were cultured overnight in Jensen’s medium. Freshly plasma bonded microfluidic flow cells were filled with sterile MJMM medium; the bubble trap was filled almost completely with medium, leaving a small air gap, and then sealed at the top with a piece of tape. The overnight culture was diluted 1:10 to OD_500_ = approx. 0.5 in MJMM and injected into the chamber via the outlet, adding just enough culture volume to fill the growth chamber. Cells were allowed to attach with no flow for 2-3 hours. This attachment OD and time are higher than those often used for flow cell biofilms of *P. aeruginosa* PAO1, but we found that they were necessary to promote robust attachment of PA14.

During this attachment period, the tubing was prepared. All tubing components were stored in 70% ethanol when not in use. Valves were used in the inlet and outlet lines so that the chamber could easily be sealed if needed. We used Tygon microbore tubing and Luer-compatible 2- and 3-way valves. For the connections to the microfluidic device, an 18-gauge blunt needle was used, with the Luer adapter removed and two 45° bends added. Inlet and outlet tubing was assembled and filled with 70% ethanol; tubing was left sitting with 70% ethanol for at least 15 minutes to ensure sterility. The tubing was then flushed extensively with sterile water and then with sterile MJMM medium. Both tubing elements were connected to the microfluidic flow cell, inlet tubing was connected to a 50 mL syringe containing fresh sterile MJMM medium, and the outlet tubing was connected to a waste bottle.

The syringe was mounted on a syringe pump, and flow was started at a rate of 10 µL/min. Given the dimensions of the growth chamber, this corresponds to an average flow speed of 400 µm/s. This flow rate means that the medium in the chamber is turning over quickly - the medium in the chamber is replaced about once a minute - but the flow is not expected to apply large mechanical forces. Consistent with this, we did not see any evidence of biofilm deformation due to flow. Biofilms were maintained under constant medium flow for the duration of the experiment, with medium-containing syringes replaced as necessary. Flow cell experiments were conducted at room temperature. When biofilms were not being imaged, they were kept covered to minimize light exposure.

### Biofilm imaging, fluorescence exclusion, and staining

Unless otherwise indicated, stains and dyes were added to the flow media at the following concentration: fluorescein: 20 µM, DiTO-1: 50 nM, Propidium Iodide (PI): 500 nM. eDNA stain. Dyes were present continuously in the media during the experiment. We use lower concentrations of DiTO-1 and PI than would be used for typical staining, but because the dye is continuously added, it accumulates over time and appropriate staining is achieved. Controls for the use of these stains and for imaging are presented in Figure S1. For single cell imaging of biofilm development, fluorescein negative staining experiments were repeated at least in biological triplicate, where a biological replicate is defined as a distinct flow cell experiment starting from a unique starter culture. Within each biological replicate, multiple biofilms were imaged across multiple fields of view. We imaged in the downstream portion of the chamber (nearer to the outlet), away from the chamber sidewalls: we consistently found that this region promoted the most robust growth of large hemispherical biofilms. For eDNA labeling and cell lysis experiments, PI and DiTO-1 labeling were used. In each case, a biofilm label was used in tandem: for PI staining, fluorescein exclusion was used, and for DiTO-1, cells constitutively expressed mApple. For mApple labeling, the fluorophore inevitably photobleached once biofilms reached a critical size. We thereafter estimated the position of the biofilm based on the linear growth observed by fluorescein exclusion imaging. To verify the final size of mApple-tagged biofilms, at the end of these experiments we stopped the flow of medium, which stopped cells from consuming oxygen and allowed the fluorophore to mature for an endpoint image. All eDNA labeling and cell lysis experiments were conducted in at least biological duplicate, with at least one replicate using fluorescein and PI and one replicate using mApple and DiTO-1. In all cases, results did not depend on which labeling method was used.

### Microscopy

All images were acquired using an Andor Dragonfly spinning disk confocal microscope. The microscope was equipped with a Nikon TI2 body, an Andor Dragonfly confocal unit, a 4.2-megapixel Zyla sCMOS camera, an Andor Integrated Laser Engine (ILE), and a motorized stage with piezo Z-axis control (ASI). A 60x 1.42 NA oil immersion objective was used for acquisition. Fluorescein and DiTO-1 images were acquired using 488 nm laser excitation and a 521/38 nm emission filter; mApple and PI images were acquired using 561 nm laser excitation and a 594/43 nm emission filter. Imaging conditions were optimized to minimize total light dosage as much as possible without compromising the information content of the images. Z stack step size was 290 nm, which provided sufficient Z sampling while mitigating total light dosage: because we were detecting cell-scale events, and the diameter of *P. aeruginosa* cells is approximately 700 nm, 290 nm Z steps provide adequate sampling. For long-duration time lapses Z stacks were acquired every 4 hours; we found that this provided adequate time resolution to monitor biofilm development while minimizing light dosage. To monitor faster time-scale events, 30-minute or 1-hour intervals were used instead. For phase contrast imaging of twitching motility, images were acquired every 2 minutes on a Nikon Ti2 eclipse, and for Airyscan, confocal Z stacks were acquired on a Zeiss LSM 980 with Airyscan 2.

### Image processing and data analysis

All image processing and data analysis was performed using custom MATLAB software. 3D registration was applied to all time lapses to correct for positional drift over time. Raw 4D data files were very large and presented a data management challenge: when possible, we downsampled images in size and bit depth, cropped regions of interest, and applied compression for easier data handling. Bit depth downsampling was performed to maximize dynamic range without introducing saturation. These transformations were applied as part of an automatic pre-processing pipeline. For display of XZ slices, an intensity correction was applied to partially compensate for signal decrease in the Z dimension due to increasing scattering. For fluorescein exclusion time lapses, intensity normalization was also applied across frames. To detect cell growth and movement, the difference was computed between adjacent time lapse frames: this difference mask has high values for dynamic regions of the image. The difference mask was overlaid on the latter of the two time frames to localize dynamics within the biofilm. For segmentation, biofilms were denoised by gaussian filtering and a binary threshold was chosen automatically by iterative selection, verified manually, and applied. For segmentation of mApple-tagged biofilms, segmentation was performed on the available frames until the fluorophore bleached; the final position was verified by endpoint imaging as described above, and the biofilm position in the intermediate frames was interpolated based on an assumed linear radial growth rate. Biofilm radius was measured as the mean distance from the biofilm surface its core.

Cell lysis events were detected as follows. First, 3D peak finding was used to find eDNA foci in each frame; peaks were filtered with an intensity cutoff that was scaled based on depth. Next, peaks were tracked over time with a linking distance of 2 µm, and tracks were filtered based on minimum length such that tracks that did not continue for the duration of the time lapse were excluded. Remaining tracks represented lysis events, with the track starting position and time indicating the lysis event. Select data sets were also analyzed manually using a custom user interface to verify the performance of the automatic lysis detection, and automatic tracking results were refined manually as needed. To estimate lysis frequency in the biofilm, because cells are extremely densely packed in the biofilm, we divided the final segmented biofilm volume into regions the size of an average *P. aeruginosa* cell^62^ and compared this to the number of lysis events to obtain a rough approximation. To compute the nearest-neighbor distances between cell lysis events, we again chose a relatively flat biofilm for simplicity and computed the XY distance from each lysis event to its nearest neighbor. We compared this to a simulated null distribution, in which we simulated the same number of points randomly distributed in the same area and measured their nearest neighbor distribution.

To visualize cell lysis events on an overlay of biofilm height (Figure 1E), we chose a relatively flat biofilm for simplicity and computed the average height (average distance from the coverslip to the biofilm surface). We then overlaid the locations of cell lysis events. Note that average biofilm height is shown; lysis events above this are from regions of the biofilm whose height is greater than the average. To measure lysis depth, we measured the distance from the lysis event to the nearest point on the biofilm surface at the time of lysis. To measure eDNA concentration vs. depth in the biofilm, we analyzed XY slices individually due to the effects of scatter on the Z intensity profile; mean eDNA staining intensity was calculated as a function of distance from the biofilm perimeter, and we used images of biofilms that had not been subjected to time lapse imaging to avoid the effects of photobleaching.

### Simulations and modeling

All simulations and models were constructed using custom MATLAB software. To create null distributions in which cell lysis occurs throughout the biofilm with equal probability, we simulated the same number of lysis events at each time point as in the underlying data set, placed them randomly within the biofilm volume at that time point, and measured the depth of each point. We repeated this 1000 times to increase the accuracy of the final prediction. To model uniform and distributed lysis, we both performed stochastic simulations and constructed models. Simulations were performed for 1- or 2D biofilms; results do not depend on the number of spatial dimensions modeled. We set the biofilm height h to increase linearly with time. For patterned lysis, simulated lysis probability was normally distributed with µ = h-5, σ = 3, and for uniform lysis, simulated lysis occurred uniformly along h. For modeling, a 1D biofilm model was used. The relationship of total cell lysis with biofilm volume was modeled as follows. For patterned lysis, in 1D the number of lysis events at each time point is constant (the area under the normal distribution), so cumulative lysis N_L_ at time t is *N_L_*(*t*)∼*t*, and ℎ(*t*)∼*t*, so *N*_*L*_(*h*)∼*h*. For uniform lysis, 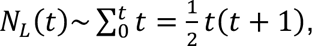 so *N*_*L*_(*h*)∼*h*(*h* + 1). The relationship of eDNA concentration [eDNA] with depth d was defined as follows. For patterned lysis, the distribution was the normal cumulative distribution function 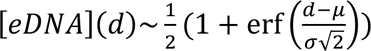 where erf is the error function and with µ = 5 and σ = 3. For uniform lysis, [*eDNA*](*d*)∼*d*. Models were fit to data with a single scaling parameter, and the model that minimized the residual sum of squares (*i.e.* maximized R^2^) was selected as best fit.

## References

1. Lyons, N.A., and Kolter, R. (2015). On the evolution of bacterial multicellularity. Curr Opin Microbiol 24, 21–28. 10.1016/j.mib.2014.12.007.

2. Kaiser, D. (2001). Building a multicellular organism. Annual Review of Genetics 35, 103–123. 10.1146/annurev.genet.35.102401.090145.

3. Bich, L., Pradeu, T., and Moreau, J.-F. (2019). Understanding multicellularity: the functional organization of the intercellular space. Front. Physiol. 10. 10.3389/fphys.2019.01170.

4. Sanz-Ezquerro, J.J., Münsterberg, A.E., and Stricker, S. (2017). Editorial: signaling pathways in embryonic development. Front. Cell Dev. Biol. 5. 10.3389/fcell.2017.00076.

5. Reuten, R., Mayorca-Guiliani, A.E., and Erler, J.T. (2022). Matritecture: mapping the extracellular matrix architecture during health and disease. Matrix Biology Plus 14, 100102. 10.1016/j.mbplus.2022.100102.

6. Hall-Stoodley, L., Costerton, J.W., and Stoodley, P. (2004). Bacterial biofilms: from the Natural environment to infectious diseases. Nat Rev Microbiol 2, 95–108. 10.1038/nrmicro821.

7. Flemming, H.-C., and Wuertz, S. (2019). Bacteria and archaea on Earth and their abundance in biofilms. Nat Rev Microbiol 17, 247–260. 10.1038/s41579-019-0158-9.

8. Vlamakis, H., Aguilar, C., Losick, R., and Kolter, R. (2008). Control of cell fate by the formation of an architecturally complex bacterial community. Genes Dev. 22, 945–953. 10.1101/gad.1645008.

9. Parsek, M.R., and Tolker-Nielsen, T. (2008). Pattern formation in *Pseudomonas aeruginosa* biofilms. Current Opinion in Microbiology 11, 560–566. 10.1016/j.mib.2008.09.015.

10. Wang, T., Shen, P., He, Y., Zhang, Y., and Liu, J. (2023). Spatial transcriptome uncovers rich coordination of metabolism in *E. coli* K12 biofilm. Nat Chem Biol 19, 940–950. 10.1038/s41589-023-01282-w.

11. Dar, D., Dar, N., Cai, L., and Newman, D.K. (2021). Spatial transcriptomics of planktonic and sessile bacterial populations at single-cell resolution. Science 373, eabi4882. 10.1126/science.abi4882.

12. Squyres, G.R., and Newman, D.K. (2024). Biofilms as more than the sum of their parts: lessons from developmental biology. Current Opinion in Microbiology 82, 102537. 10.1016/j.mib.2024.102537.

13. Walma, D.A.C., and Yamada, K.M. (2020). The extracellular matrix in development. Development 147, dev175596. 10.1242/dev.175596.

14. Flemming, H.-C., and Wingender, J. (2010). The biofilm matrix. Nat Rev Microbiol 8, 623–633. 10.1038/nrmicro2415.

15. Flemming, H.-C., van Hullebusch, E.D., Neu, T.R., Nielsen, P.H., Seviour, T., Stoodley, P., Wingender, J., and Wuertz, S. (2023). The biofilm matrix: multitasking in a shared space. Nat Rev Microbiol 21, 70–86. 10.1038/s41579-022-00791-0.

16. Whitchurch, C.B., Tolker-Nielsen, T., Ragas, P.C., and Mattick, J.S. (2002). Extracellular DNA required for bacterial biofilm formation. Science 295, 1487. 10.1126/science.295.5559.1487.

17. Okshevsky, M., and Meyer, R.L. (2015). The role of extracellular DNA in the establishment, maintenance and perpetuation of bacterial biofilms. Crit Rev Microbiol 41, 341–352. 10.3109/1040841X.2013.841639.

18. Okshevsky, M., Regina, V.R., and Meyer, R.L. (2015). Extracellular DNA as a target for biofilm control. Current Opinion in Biotechnology 33, 73–80. 10.1016/j.copbio.2014.12.002.

19. Allesen-Holm, M., Barken, K.B., Yang, L., Klausen, M., Webb, J.S., Kjelleberg, S., Molin, S., Givskov, M., and Tolker-Nielsen, T. (2006). A characterization of DNA release in *Pseudomonas aeruginosa* cultures and biofilms. Mol Microbiol 59, 1114–1128. 10.1111/j.1365-2958.2005.05008.x.

20. Jennings, L.K., Storek, K.M., Ledvina, H.E., Coulon, C., Marmont, L.S., Sadovskaya, I., Secor, P.R., Tseng, B.S., Scian, M., Filloux, A., et al. (2015). Pel is a cationic exopolysaccharide that cross-links extracellular DNA in the *Pseudomonas aeruginosa* biofilm matrix. Proceedings of the National Academy of Sciences 112, 11353–11358. 10.1073/pnas.1503058112.

21. Turnbull, L., Toyofuku, M., Hynen, A.L., Kurosawa, M., Pessi, G., Petty, N.K., Osvath, S.R., Cárcamo-Oyarce, G., Gloag, E.S., Shimoni, R., et al. (2016). Explosive cell lysis as a mechanism for the biogenesis of bacterial membrane vesicles and biofilms. Nat Commun 7, 11220. 10.1038/ncomms11220.

22. Prentice, J.A., Kasivisweswaran, S., Weerd, R. van de, and Bridges, A.A. (2024). Biofilm dispersal patterns revealed using far-red fluorogenic probes. Preprint at bioRxiv, 10.1101/2024.07.15.603607 10.1101/2024.07.15.603607.

23. Hynen, A.L., Lazenby, J.J., Savva, G.M., McCaughey, L.C., Turnbull, L., Nolan, L.M., and Whitchurch, C.B. (2021). Multiple holins contribute to extracellular DNA release in *Pseudomonas aeruginosa* biofilms. Microbiology (Reading) 167, 000990. 10.1099/mic.0.000990.

24. Drescher, K., Dunkel, J., Nadell, C.D., van Teeffelen, S., Grnja, I., Wingreen, N.S., Stone, H.A., and Bassler, B.L. (2016). Architectural transitions in *Vibrio cholerae* biofilms at single-cell resolution. Proceedings of the National Academy of Sciences 113, E2066–E2072. 10.1073/pnas.1601702113.

25. Tolker-Nielsen, T., Brinch, U.C., Ragas, P.C., Andersen, J.B., Jacobsen, C.S., and Molin, S. (2000). Development and dynamics of *Pseudomonas*sp. biofilms. Journal of Bacteriology 182, 6482–6489. 10.1128/jb.182.22.6482-6489.2000.

26. Yan, J., Sharo, A.G., Stone, H.A., Wingreen, N.S., and Bassler, B.L. (2016). *Vibrio cholerae* biofilm growth program and architecture revealed by single-cell live imaging. Proceedings of the National Academy of Sciences 113, E5337–E5343. 10.1073/pnas.1611494113.

27. Jelli, E., Ohmura, T., Netter, N., Abt, M., Jiménez-Siebert, E., Neuhaus, K., Rode, D.K.H., Nadell, C.D., and Drescher, K. (2023). Single-cell segmentation in bacterial biofilms with an optimized deep learning method enables tracking of cell lineages and measurements of growth rates. Molecular Microbiology 119, 659–676. 10.1111/mmi.15064.

28. Stewart, P.S., Zhang, T., Xu, R., Pitts, B., Walters, M.C., Roe, F., Kikhney, J., and Moter, A. (2016). Reaction–diffusion theory explains hypoxia and heterogeneous growth within microbial biofilms associated with chronic infections. npj Biofilms Microbiomes 2, 1–8. 10.1038/npjbiofilms.2016.12.

29. Caldwell, D.E., Korber, D.R., and Lawrence, J.R. (1992). Imaging of bacterial cells by fluorescence exclusion using scanning confocal laser microscopy. Journal of Microbiological Methods 15, 249–261. 10.1016/0167-7012(92)90045-6.

30. Smart, P.L., and Laidlaw, I.M.S. (1977). An evaluation of some fluorescent dyes for water tracing. Water Resources Research 13, 15–33. 10.1029/WR013i001p00015.

31. Purevdorj-Gage, B., Costerton, W.J., and Stoodley, P. (2005). Phenotypic differentiation and seeding dispersal in non-mucoid and mucoid *Pseudomonas aeruginosa* biofilms. Microbiology 151, 1569–1576. 10.1099/mic.0.27536-0.

32. Barraud, N., Kjelleberg, S., and Rice, S.A. (2015). Dispersal from microbial biofilms. Microbiology Spectrum 3, 10.1128/microbiolspec.mb-0015–2014. 10.1128/microbiolspec.mb-0015-2014.

33. Ma, L., Conover, M., Lu, H., Parsek, M.R., Bayles, K., and Wozniak, D.J. (2009). Assembly and development of the *Pseudomonas aeruginosa* biofilm matrix. PLOS Pathogens 5, e1000354. 10.1371/journal.ppat.1000354.

34. Webb, J.S., Thompson, L.S., James, S., Charlton, T., Tolker-Nielsen, T., Koch, B., Givskov, M., and Kjelleberg, S. (2003). Cell death in *Pseudomonas aeruginosa* biofilm development. J Bacteriol 185, 4585–4592. 10.1128/JB.185.15.4585-4592.2003.

35. Stewart, P.S., and Franklin, M.J. (2008). Physiological heterogeneity in biofilms. Nat Rev Microbiol 6, 199–210. 10.1038/nrmicro1838.

36. Jo, J., Price-Whelan, A., and Dietrich, L.E.P. (2022). Gradients and consequences of heterogeneity in biofilms. Nat Rev Microbiol 20, 593–607. 10.1038/s41579-022-00692-2.

37. Chou, K.-T., Lee, D.D., Chiou, J., Galera-Laporta, L., Ly, S., Garcia-Ojalvo, J., and Süel, G.M. (2022). A segmentation clock patterns cellular differentiation in a bacterial biofilm. Cell 185, 145–157.e13. 10.1016/j.cell.2021.12.001.

38. Stevanovic, M., Boukéké-Lesplulier, T., Hupe, L., Hasty, J., Bittihn, P., and Schultz, D. (2022). Nutrient gradients mediate complex colony-level antibiotic responses in structured microbial populations. Front. Microbiol. 13. 10.3389/fmicb.2022.740259.

39. Meirelles, L.A., and Newman, D.K. (2018). Both toxic and beneficial effects of pyocyanin contribute to the lifecycle of *Pseudomonas aeruginosa*. Molecular Microbiology 110, 995–1010. 10.1111/mmi.14132.

40. Lenne, P.-F., Munro, E., Heemskerk, I., Warmflash, A., Bocanegra-Moreno, L., Kishi, K., Kicheva, A., Long, Y., Fruleux, A., Boudaoud, A., et al. (2021). Roadmap for the multiscale coupling of biochemical and mechanical signals during development. Phys Biol 18, 10.1088/1478-3975/abd0db. 10.1088/1478-3975/abd0db.

41. Suzanne, M., and Steller, H. (2013). Shaping organisms with apoptosis. Cell Death Differ 20, 669–675. 10.1038/cdd.2013.11.

42. Li, M., Nahum, Y., Matouš, K., Stoodley, P., and Nerenberg, R. (2023). Effects of biofilm heterogeneity on the apparent mechanical properties obtained by shear rheometry. Biotechnology and Bioengineering 120, 553–561. 10.1002/bit.28276.

43. Otto, S.B., Martin, M., Schäfer, D., Hartmann, R., Drescher, K., Brix, S., Dragoš, A., and Kovács, Á.T. (2020). Privatization of biofilm matrix in structurally heterogeneous biofilms. mSystems 5, e00425–20. 10.1128/mSystems.00425-20.

44. Saunders, S.H., Tse, E.C.M., Yates, M.D., Otero, F.J., Trammell, S.A., Stemp, E.D.A., Barton, J.K., Tender, L.M., and Newman, D.K. (2020). Extracellular DNA promotes efficient extracellular electron transfer by pyocyanin in *Pseudomonas aeruginosa* biofilms. Cell 182, 919–932.e19. 10.1016/j.cell.2020.07.006.

45. Bellin, D.L., Sakhtah, H., Zhang, Y., Price-Whelan, A., Dietrich, L.E.P., and Shepard, K.L. (2016). Electrochemical camera chip for simultaneous imaging of multiple metabolites in biofilms. Nat Commun 7, 10535. 10.1038/ncomms10535.

46. Häussler, S., and Becker, T. (2008). The pseudomonas quinolone signal (PQS) balances life and death in *Pseudomonas aeruginosa* populations. PLOS Pathogens 4, e1000166. 10.1371/journal.ppat.1000166.

47. Whiteley, M., Lee, K.M., and Greenberg, E.P. (1999). Identification of genes controlled by quorum sensing in *Pseudomonas aeruginosa*. Proceedings of the National Academy of Sciences 96, 13904–13909. 10.1073/pnas.96.24.13904.

48. Kolter, R., and Greenberg, E.P. (2006). The superficial life of microbes. Nature 441, 300–302. 10.1038/441300a.

49. van Gestel, J., Vlamakis, H., and Kolter, R. (2015). Division of labor in biofilms: the ecology of cell differentiation. Microbiology Spectrum 3, 10.1128/microbiolspec.mb-0002–2014. 10.1128/microbiolspec.mb-0002-2014.

50. Shilo, B.-Z., and Barkai, N. (2017). Buffering global variability of morphogen gradients. Developmental Cell 40, 429–438. 10.1016/j.devcel.2016.12.012.

51. Grieneisen, V.A., Xu, J., Marée, A.F.M., Hogeweg, P., and Scheres, B. (2007). Auxin transport is sufficient to generate a maximum and gradient guiding root growth. Nature 449, 1008–1013. 10.1038/nature06215.

52. Klausen, M., Heydorn, A., Ragas, P., Lambertsen, L., Aaes-Jørgensen, A., Molin, S., and Tolker-Nielsen, T. (2003). Biofilm formation by *Pseudomonas aeruginosa* wild type, flagella and type IV pili mutants. Molecular Microbiology 48, 1511–1524. 10.1046/j.1365-2958.2003.03525.x.

53. Hunt, S.M., Werner, E.M., Huang, B., Hamilton, M.A., and Stewart, P.S. (2004). Hypothesis for the role of nutrient starvation in biofilm detachment. Appl Environ Microbiol 70, 7418–7425. 10.1128/AEM.70.12.7418-7425.2004.

54. Sauer, K., Cullen, M.C., Rickard, A.H., Zeef, L.A.H., Davies, D.G., and Gilbert, P. (2004). Characterization of nutrient-induced dispersion in *Pseudomonas aeruginosa* PAO1 biofilm. J Bacteriol 186, 7312–7326. 10.1128/JB.186.21.7312-7326.2004.

55. Stoodley, P., Sauer, K., Davies, D.G., and Costerton, J.W. (2002). Biofilms as complex differentiated communities. Annual Review of Microbiology 56, 187–209. 10.1146/annurev.micro.56.012302.160705.

56. Shanks, R.M.Q., Caiazza, N.C., Hinsa, S.M., Toutain, C.M., and O’Toole, G.A. (2006). *Saccharomyces cerevisiae*-based molecular tool kit for manipulation of genes from gram-negative bacteria. Appl Environ Microbiol 72, 5027–5036. 10.1128/AEM.00682-06.

57. Lambertsen, L., Sternberg, C., and Molin, S. (2004). Mini-Tn7 transposons for site-specific tagging of bacteria with fluorescent proteins. Environmental Microbiology 6, 726–732. 10.1111/j.1462-2920.2004.00605.x.

58. Lanzer, M., and Bujard, H. (1988). Promoters largely determine the efficiency of repressor action. Proceedings of the National Academy of Sciences 85, 8973–8977. 10.1073/pnas.85.23.8973.

59. Hoang, T.T., Kutchma, A.J., Becher, A., and Schweizer, H.P. (2000). Integration-proficient plasmids for *Pseudomonas aeruginosa:* site-specific integration and use for engineering of reporter and expression strains. Plasmid 43, 59–72. 10.1006/plas.1999.1441.

60. Choi, K.-H., and Schweizer, H.P. (2006). mini-Tn7 insertion in bacteria with single attTn7 sites: example *Pseudomonas aeruginosa*. Nat Protoc 1, 153–161. 10.1038/nprot.2006.24.

61. Cont, A., Rossy, T., Al-Mayyah, Z., and Persat, A. (2020). Biofilms deform soft surfaces and disrupt epithelia. eLife 9, e56533. 10.7554/eLife.56533.

62. Iglewski, B.H. (1996). Pseudomonas. In Medical Microbiology, S. Baron, ed. (University of Texas Medical Branch at Galveston).

