## Supplementary material for "Real-time high-resolution microscopy reveals how single-cell lysis shapes biofilm matrix morphogenesis": Figures S1-S3 + Supp Video legends

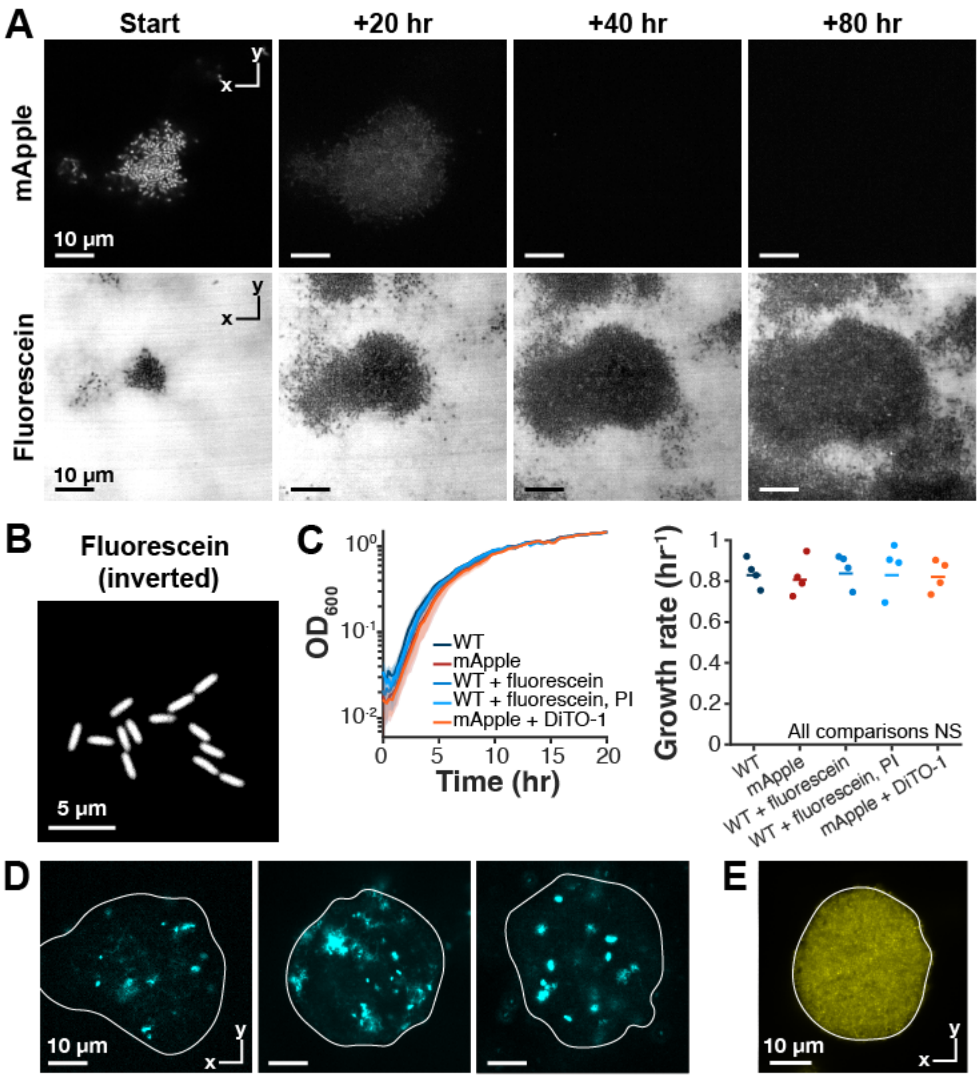


**Figure S1: Fluorescence exclusion microscopy and eDNA staining**

**(A)** Fluorescence exclusion microscopy enables long-term imaging of biofilm development. Comparison of biofilm labeling with cells expressing mApple vs. fluorescence exclusion imaging. Fluorescein exclusion allows live biofilm imaging with no loss in performance over time, whereas fluorescent protein signal is lost in mature biofilms. XY confocal slices of 4D biofilm imaging. **(B)** Fluorescence exclusion enables high-quality single cell imaging. Image is shown inverted for comparison to conventional fluorescent protein imaging. XY confocal slice of single cells. **(C)** The dyes used in this study and mApple fluorescent protein expression do not impact cell growth. Left: growth curves of the indicated strains and dye combinations: these are the same strains and dyes in the same combinations and at the same concentrations used throughout the study. Lines: mean of 4 biological replicates, shading: standard deviation. Right: growth rates obtained by fitting the growth curves at left. Differences between growth rates are not significant by T-test. **(D)** Staining and time-lapse imaging do not strongly alter eDNA architecture. Representative XY confocal slices of eDNA stained with DiTO-1 (cyan), white line indicates biofilm surface. Left: biofilm grown in the presence of 50 nM DiTO-1 and imaged for 104 hours, center: biofilm grown in the presence of 50 nM DiTO-1 without time lapse imaging, and right: biofilm grown without staining or imaging, stained with 1 μM DiTO-1 immediately prior to imaging. Other than photobleaching decreasing the fluorescence signal at left, no strong effects are detected from growth in the presence of DiTO-1 or time lapse imaging. **(E)** eDNA stains can label the entire biofilm. Biofilm grown in medium without carbon for 3 days, leading to complete labeling with PI (yellow). White line indicates biofilm surface. The resulting PI stain is distributed evenly throughout the biofilm, indicating that observed patterns in eDNA staining are not due to biases in labeling.

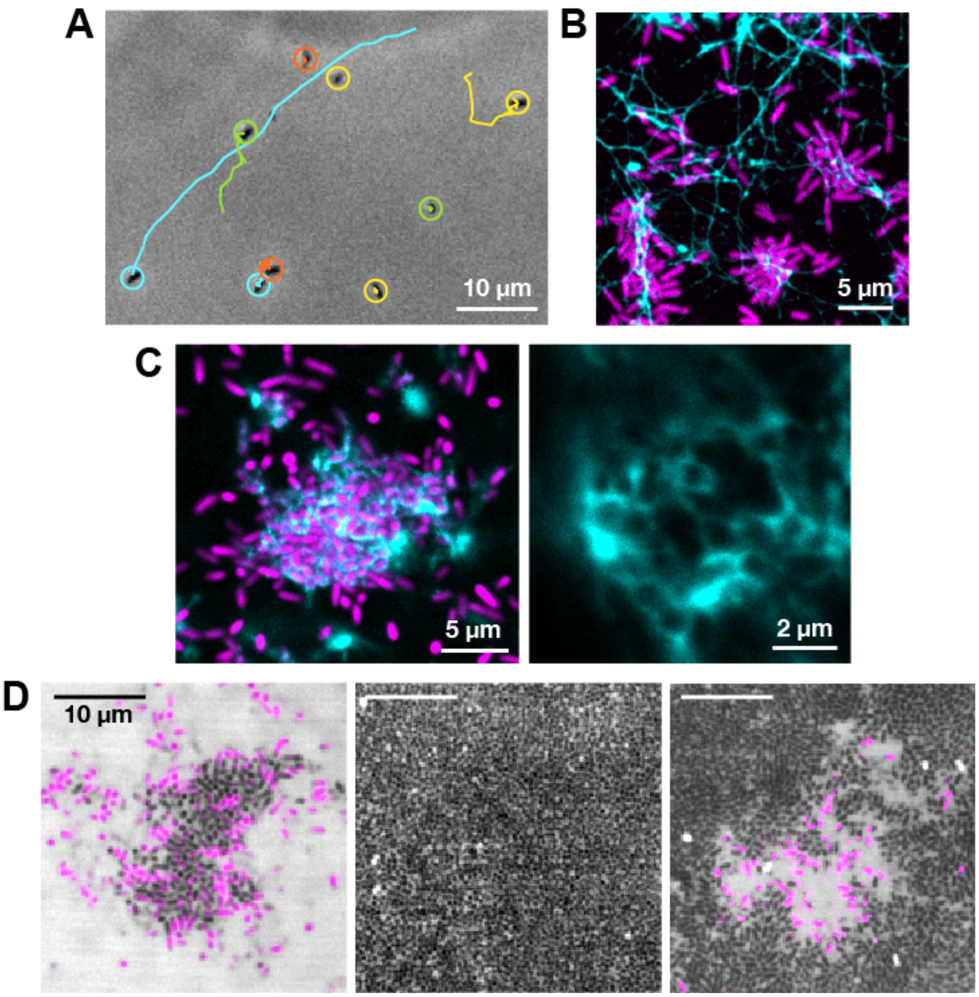


**Figure S2: Supplemental biofilm imaging**

**(A)** In the attachment phase, cells are sparse and exhibit twitching motility. Cells were imaged during the attachment phase by phase contrast microscopy every 2 minutes for 50 minutes. Cell movement was tracked: cells are circled, with their tracks shown as lines. Some cells are stationary, but others exhibit twitching movement. **(B)** As microcolonies begin to form, eDNA is present in a web on the surface. Microcolonies contain eDNA, as expected[23]. Cells labeled with mApple (magenta), eDNA labeled with DiTO-1 (cyan). **(C)** In microcolonies, cells are individually encircled by eDNA. Left: XY confocal section, cells labeled with mApple (magenta), eDNA labeled with DiTO-1 (cyan). Right: Airyscan section of eDNA architecture in microcolonies. **(D)** Cell organization and dynamics over time. XY confocal sections of the core of the biofilm during microcolony (left), mature biofilm (center), and dispersal stages (right), visualized by fluorescein exclusion. Cells colored magenta are dynamic, moving and/or growing during the previous 4 hours. In microcolonies, cells are more loosely packed and dynamic throughout. In the core of mature biofilms, cells are densely packed and do not detectably move or grow. During dispersal, a small number of cells in the biofilm core resume their motility.

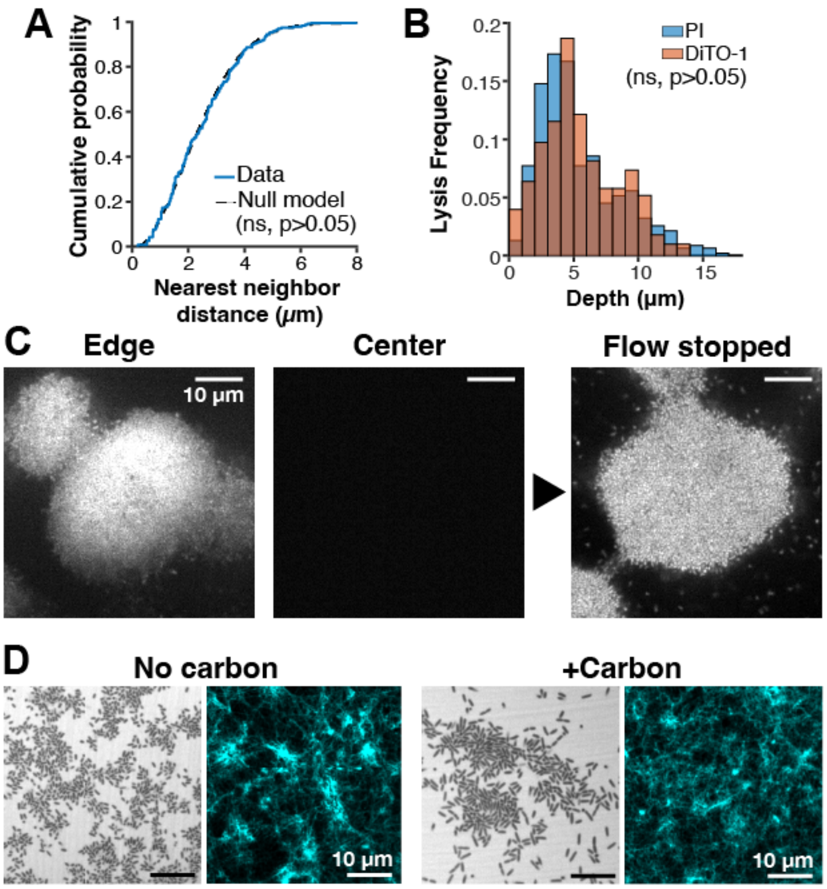


**Figure S3: Controls for cell lysis patterning studies**

**(A)** Cell lysis is stochastic on small length scales. Cell lysis is patterned on the biofilm scale, but within the lysis zone, the nearest neighbor distances match a simulated random distribution (not significantly different by Mann-Whitney U test). **(B)** The cell lysis distribution does not depend on the eDNA stain used. Comparison of lysis distributions measured with PI (blue) and DiTO-1 (red); distributions are not significantly different by Mann-Whitney U test. **(C)** The microfluidic device contains an oxygen-rich region near the chamber edge. After 70 hours of time-lapse imaging, biofilms in the center no longer have detectable mApple fluorescence, but edge biofilms do. This suggests that biofilms in the bulk of the flow cell have lower oxygen, preventing fluorescent protein maturation, but that biofilms at the edge have more oxygen available. To confirm this, we stopped the flow: oxygen levels increase throughout the flow cell (because cellular consumption of oxygen slows down), and fluorescence indeed recovers in the non-edge biofilm because of fluorophore maturation. **(D)** Removing carbon from the medium does not lead to dysregulation of cell lysis in the absence of biofilm formation. Cells in the attachment phase were grown with or without carbon in the medium for 3 days, with no clear impact on cells (fluorescein exclusion, left) or eDNA (DiTO-1 stain, cyan, right). XY confocal slices.

**Supplemental video legends:**

**Video S1: Biofilm growth by fluorescein exclusion imaging,** companion to Figure 1C. XZ confocal section of time lapse of biofilm growth. Note that cell permeability to fluorescein is also depth-dependent in time, with a similar depth profile to the cell lysis distribution we observe. Video duration: 108 hours, Scale bar, 5 μm.

**Video S2: The biofilm core lifts during dispersal,** companion to Figures 1D and 2B. Confocal sections of time lapses of hollow core formation during biofilm dispersal. Top left: XZ section of fluorescein exclusion imaging; top right: XZ section of DiTO-1 labeling, bottom: XY section of DiTO-1 labeling. Video duration: 36 hours, Scale bar, 5 μm.

**Video S3: Patterning of cell lysis during biofilm growth**, companion to Figure 2D. XZ confocal section of time lapse of cell lysis during biofilm growth, labeled with PI. White line: biofilm surface, segmented based on fluorescein exclusion imaging. Video duration: 100 hours, Scale bar, 5 μm.

**Video S4: Cell death patterning is lost in the absence of nutrient gradients,** companion to Figure 4E. XZ confocal section of time lapse of cell lysis during biofilm growth; carbon was removed from the medium at the start of the time lapse. Video duration: 36 hours, Scale bar, 5 μm.

**Video S5: Cell death occurs in the biofilm core when oxygen is high,** companion to Figure 4H. XZ confocal section of time lapse of cell lysis during biofilm growth in a high-oxygen region of the flow cell, labeled with PI. Video duration: 108 hours, Scale bar, 5 μm.
